# Condensation of the RNA chaperone Hfq is coupled to inhibition of carbon assimilation and contributes to the stabilisation of regulatory RNAs in nitrogen starved *Escherichia coli*

**DOI:** 10.1101/2025.05.19.654817

**Authors:** Josh McQuail, Harriet Ellis, Volker Behrends, Cristina Balcells-Nadal, Thorsten Bischler, Tom Gräfenhan, Sivaramesh Wigneshweraraj

## Abstract

Ribonucleoprotein-condensates are membraneless compartments that concentrate RNA-binding proteins and RNA and play key roles in cellular adaptation across both eukaryotes and bacteria. While the biological roles of ribonucleoprotein-condensates are better understood in eukaryotic systems, the knowledge of metabolic processes that govern their formation and their contribution to stress adaptation remains at a nascent stage in bacterial RNA biology. Hfq is an RNA-chaperone conserved in many bacteria that undergoes condensation in response to diverse stresses. Using nitrogen (N) starvation in *Escherichia coli* as a model stress condition, we show that Hfq condensation occurs independently of any extracellular cues, cytoplasmic shrinkage that cells undergo during N starvation or the canonical NtrBC-dependent adaptive response to N starvation. However, we demonstrate that Hfq condensation is coupled to the inhibition of carbon assimilation in N-starved *E. coli*. Further, by comparing the transcriptomes of wild-type bacteria and bacteria unable to form Hfq-condensates, we reveal that Hfq-condensates contribute to the stabilisation of Hfq-associated non-coding regulatory RNAs. We propose that coordination of carbon and N metabolism during N starvation, critical for metabolic adaptation, is accompanied by preservation of non-coding regulatory RNAs via Hfq condensation.

## Introduction

Bacteria often exist in a growth attenuated state for long periods of time, as many environments inhabited by bacteria are limited for nutrients. As such, bacteria exhibit a range of adaptive responses to long-term nutrient starvation. Many of these are underpinned by changes in gene expression at the level of RNA metabolism. At the level of RNA synthesis, specific transcription regulators respond to cellular cues and modulate the activity of the RNA polymerase, reprogramming the transcriptional landscape and thereby activating or downregulating specific metabolic pathways. In parallel, regulatory non-coding RNA molecules, hereafter referred to as small RNAs (sRNA), mediate the post-transcriptional regulation of gene expression in bacteria. Some sRNAs are *cis*-acting and are expressed from the same locus as their sole mRNA target. However, the majority of sRNA are *trans*-acting and target multiple mRNA targets at different genetic loci. As such, unlike *cis*-acting sRNAs, *trans*-acting sRNAs share only partial complementarity with their mRNA targets and significantly depend on RNA binding chaperone proteins to interact with their targets. sRNA can affect the fate of their cognate mRNA in at least four different ways (reviewed in (1–3)): (i) they facilitate translation of the target mRNA by altering RNA secondary structure and allowing ribosomes access to the Shine-Dalgarno sequence; (ii) they prevent ribosomes access the Shine-Dalgarno sequence, translationally repressing the target mRNA, (iii) they enable the rapid degradation of target mRNA by the RNA degradosome or (iv) they protect the target mRNA from being degraded by RNases.

Nitrogen (N) is used for the biosynthesis of the building blocks of all proteins (amino acids), nucleic acids (nucleotides) as well as numerous metabolites and cofactors in bacteria. As such, N is an essential component of the bacterial cell. Many groups have used *Escherichia coli* as the model bacterium to understand the adaptive response to N-starvation - the transcriptional regulatory basis of the adaptive-response to N-starvation is well-understood (4). Briefly, this involves the two-component system, NtrBC, in which the phosphorylation of the response regulator NtrC by its cognate sensor kinase NtrB allows NtrC-P to activate the transcription of ∼100 genes. These include *relA*, the gene responsible for the synthesis of the RNA polymerase binding stress signalling nucleotide guanosine pentaphosphate (5), and *nac*, which encodes a global transcription regulator - nitrogen assimilation control protein, Nac (6). Consequently, this leads to a large-scale reprogramming of the transcriptome to allow N scavenging from alternative sources of N, concomitant with the quiescing of cellular processes which consume N (5,7,8). In contrast, very little is known about the post-transcriptional regulatory basis of the adaptive response to N starvation in bacteria. Emerging results have now begun to shed light on the role of Hfq-dependent sRNAs in N-starved *E. coli*. Walling et al. showed that the sRNA GlnZ promotes cell survival by regulating genes linked to nitrogen and carbon flux in N starved *E. coli* (see later) (9). McQuail et al. used genome-scale analyses to reveal that Hfq-mediated sRNA-mRNA interactions in *E. coli* are extensive and dynamic during N starvation (10), indicating a continuous requirement for post-transcriptional regulation during N-starvation.

Notably, we previously showed that ∼50% of all Hfq molecules gradually assemble into foci-like subcellular condensates near the cell poles as a function of time under N starvation by a process analogous to liquid-liquid phase separation (11,12). These Hfq-condensates are notably reversible, i.e. they disperse when N is replenished. Further, the Hfq-condensates co-localised with condensates of the RNA-degradosome (11). In a subsequent study, Goldberg et al. showed that Hfq-condensates also exist in stationary phase *E. coli* grown in lysogeny broth or in bacteria experiencing osmotic shock, suggesting that the Hfq-condensates are a prevalent feature in the subcellular landscape of stressed bacteria (13). The Hfq-condensates that form in stationary phase and osmotically stressed *E. coli* did not co-localise with the RNA-degradosome but with a condensate formed by a protein called TmaR. The TmaR protein binds to and inhibits the phosphotransferase system (PTS) enzyme I, and the presence of TmaR-condensates is indicative of inhibition of sugar uptake (14,15). Strikingly, the Hfq-condensates do not form in cells devoid of *tmaR*, indicating that condensation of Hfq in stationary phase and osmotically stressed bacteria depends on TmaR-condensates; conversely, the TmaR-condensates form independently of Hfq-condensates. Whether TmaR condensates are required for Hfq condensation during N starvation is unknown.

We further showed that the efficacy by which the Hfq-condensates form in N-starved *E. coli* is affected by the RNA binding activity of Hfq (12). Consistent with this view, Goldberg et al. demonstrated that RNA is a constituent of the Hfq-condensates by assembling Hfq-condensates *in vitro*. This study further showed that some polar-enriched RNA molecules in osmotically stressed bacteria, including sRNAs, were destabilised in Δ*tmaR* bacteria (which do not contain Hfq-condensates), implying, but not directly demonstrating, that the Hfq-condensates have a role in RNA stabilisation (13,16). A recent study by Guan et al. (17) revealed that long-chain polyphosphates (polyP), which accumulates during N starvation, serves as a scaffold for the assembly of the Hfq-condensates. This study also supported a role for the Hfq-condensates in RNA stabilisation and showed that Hfq-condensates are a nucleoprotein hub, serving to protect mRNA transcripts, while simultaneously facilitating the degradation of transcripts less essential for survival and growth resumption.

A characteristic property of the Hfq-condensates in N-starved *E. coli* is that they form gradually as a function of time during N starvation (12,17). Under conditions used in our previous studies (11,12) and in the recent study by Guan et al., Hfq-condensates are not readily detected until ∼6 hours following N run-out in the media. After which time, the Hfq-condensates gradually formed and, ∼80-90% of all cells contained the Hfq-condensates by 24 hours into N starvation (N-24). Interestingly, in an *E. coli* strain devoid of polyP (Δ*ppk*), no Hfq-condensates are detected following 6 hours of N starvation (17); however, ∼30-40% of cells in the Δ*ppk* population still contained Hfq-condensates 24 hours after the onset of N starvation (17). Therefore, whilst polyP certainly contributes to the formation of Hfq-condensates, it is clearly not strictly required, nor sufficient, for Hfq-condensate formation.

The formation of biomolecular condensates by RNA and proteins associated with RNA metabolism, henceforth referred to as ribonucleoprotein condensates, is an emergent and prevalent feature in the subcellular landscape of diverse stressed bacteria (reviewed in (18)). However, we are yet to understand what metabolic cellular processes drive the formation of ribonucleoprotein condensates, or how they contribute to adaptive stress responses. In the current study, we used Hfq-condensates that form under N starvation to address some of these outstanding questions.

## Materials and Methods

### Bacterial strains and plasmids

All strains used in this study were derived from *Escherichia coli* K-12 and are listed in Supplementary Table 1. The TmaR-PAmCherry strain was constructed using the λ Red recombination method (19) to create an in-frame fusion encoding a linker sequence and PAmCherry, followed by a kanamycin resistance cassette (amplified from the Hfq-PAmCherry strain) to the 3′ end of *tmaR*. Gene deletions (Δ*tmaR* or Δ*ppK*) were introduced into the Hfq-PAmCherry strain as described previously (5): briefly, the knockout alleles were transduced using the P1*vir* bacteriophage with strains from the Keio collection (20) serving as donors.

### Bacterial growth conditions

N starvation experiments were conducted as described in (8). Briefly, unless otherwise stated bacteria were grown in Gutnick minimal medium (33.8 mM KH_2_PO_4_, 77.5 mM K_2_HPO_4_, 5.74 mM K_2_SO_4_, 0.41 mM MgSO_4_), supplemented with Ho-LE trace elements (21), and 0.4% (w/v) glucose, and 10 mM NH_4_Cl (for overnight cultures) or 3 mM NH_4_Cl (for N starvation experiments) at 37°C in a shaking (180 rpm) incubator. For experiments involving spent media: cultures of WT MG1655 *E. coli* were first grown to N-24 and centrifuged, the resulting ‘spent’ media was then sterile filtered before use. Bacteria from either N+ or N-were centrifuged, resuspended in spent media and incubated at 37°C in a shaking (180 rpm) incubator. For experiments involving dimethyl-ketoglutarate (dmKG), dmKG was added directly into the culture to a final concentration of 40mM at N-. The proportion of viable cells in the bacterial population was determined by measuring CFU/ml from serial dilutions on lysogeny broth agar plates.

### Photoactivated localization microscopy (PALM) and single molecule tracking (SMT)

For the PALM and single-molecular tracking (SMT) experiments, the Hfq-PAmCherry (and derivatives) and TmaR-PAmCherry reporter strains were used. Bacterial cultures were grown as described above and samples were taken at the indicated time points, then imaged and analysed as previously described (22,23). Briefly, 1 ml of culture was centrifuged, washed and resuspended in a small amount of Gutnick minimal medium without any NH_4_Cl + 0.4% glucose; samples taken at N+ were resuspended in Gutnick minimal medium with 3 mM NH_4_Cl + 0.4% glucose; samples in C-source experiments were resuspended in Gutnick minimal media without either NH_4_Cl or glucose. One μl of the resuspended culture was then placed on a Gutnick minimal medium agarose pad (1x Gutnick minimal medium with no NH_4_Cl + 0.4% glucose with 1% (w/v) agarose); samples taken at N+ were placed on a pad made with Gutnick minimal medium with 3 mM NH_4_Cl, samples in C-source experiments were placed on a pad made with Gutnick minimal media without either NH_4_Cl or glucose. Cells were imaged on a PALM-optimized Nanoimager (Oxford Nanoimaging, https://oni.bio/nanoimager/) with 15 millisecond exposures, at 66 frames per second over 10,000 frames. Photoactivatable molecules were activated using 405 nm and 561 nm lasers. Fields-of-view typically consisted of 100-200 bacterial cells. For SMT, the Nanoimager software was used to localize the molecules by fitting detectable spots of high photon intensity to a Gaussian function. The Nanoimager software SMT function was then used to track individual molecules and draw trajectories of individual molecules over multiple frames, using a maximum step distance between frames of 0.6 μm and a nearest-neighbour exclusion radius of 0.9 μm. The software then calculated the apparent diffusion coefficients (*D\**) for each trajectory over at least four steps, based on the mean squared displacement of the molecule. To calculate %H_IM_, we collated D* values from multiple fields of view and determined the proportion of D* values that fell into our previously defined immobile population (D* Σ0.08 μm/s^2^). To calculate the proportion of cells with foci, PALM datasets were first analysed using the Cluster analysis function of CODI (Oxford Nanoimaging, https://alto.codi.bio/). Both Hfq and TmaR foci were analysed using DBSCAN*: Hfq foci were defined with an eps distance of 75nm and filtered to those with >50 localisations/cluster and a density >0.0015 localisations/nm^2^, TmaR foci were defined with an eps distance of 100nm and filtered to those with >100 localisations/cluster. Total number of foci per field of view was then used to determine the portion of total cells which contained foci.

### Glucose assay

Glucose concentration in the media was determined using the Glucose Assay Kit (Sigma-Aldrich, MAK476) in accordance with the manufacturer’s protocol.

### Targeted metabolite measurement

At the indicated time points, approximately 10^10^ cells were collected, and washed twice in ¼ strength Ringer’s solution (Thermo Scientific, BR0052G). Cell pellets were resuspended in 500μl of cold (-20°C) methanol:acetonitrile:water (2:2:1, v/v/v) + 0.1% formic acid. Samples were stored at -80°C until analysis. Before analysis, all vials received were reconstituted with 150 µL of 97.5% H_2_O + 2.5% acetonitrile + 0.2% formic acid (FA), diluted, vortexed and transferred to inserts. Pooled quality control of all samples was then generated by pooling 10 µL of the first replicate for each experimental condition and injected every 8 samples. All reagents used were of ultra-high-performance liquid chromatography (UHPLC) gradient grade and all standards were of analytical grade. Targeted metabolomics analysis was performed using an Agilent 1290 Liquid chromatography (LC) system (Agilent Technologies, CA, USA) coupled to a QTRAP 4000 mass spectrometry (MS) system (SCIEX, Danaher, WA, USA). Chromatographic separation was achieved using a Luna Omega Polar C18 column (Phenomenex/Danaher, WA, USA). The analysis was conducted in positive ion mode (A: H2O + 0.2% FA / B: acetonitrile + 0.2% FA) on a 20-minute gradient and in negative ion mode (A: H2O + 0.1% FA + 10 mM ammonium formate / B: 100% acetonitrile) on a 14-minute gradient at 0.450 ml/min flow rate. All data was acquired in multiple reaction monitoring (MRM) mode. Resulting spectra analysed using an in-house data analysis workflow based on (24).

### T7 phage infection assay

Bacterial cultures were grown in Gutnick minimal medium as described above to the indicated time points. Bacterial culture samples were taken, centrifuged and resuspended in fresh Gutnick minimal media supplemented with either 2mM NH_4_Cl and 12.5mM glucose (for N+) or 5mM glucose (for N-24) and diluted to *A*_600 nm_ of 0.3 to a final volume of 500 μl and transferred to a flat-bottomed 48-well plate, together with T7 phage at a final concentration of 4.2 × 10^9^ phage/ml. The cultures were then grown at 37 °C with shaking at 700 rpm in a SPECTROstar Nano microplate reader (BMG LABTECH), and *A*_600 nm_ readings were taken every 10 min.

### RNA sequencing

WT, Δ*tmaR* and Δ*hfq* bacteria were harvested at N-24; all strains contained the empty plasmid pBAD18, as this was performed alongside a larger transcriptomics project. RNA was extracted using the RNAsnap protocol (25). Three biological replicates of each strain were taken and mixed with a phenol:ethanol (1:19) solution at a ratio of 9:1 (culture:solution) before harvesting the bacteria immediately by centrifugation. Pellets were resuspended in RNA extraction solution (18 mM EDTA, 0.025% SDS, 1% 2-mercaptoethanol, 95% formamide) and lysed at 95°C for 10 min. Cell debris was pelleted by centrifugation. RNA was purified with PureLink RNA Mini Kit extraction columns (Invitrogen, 12183018A) and largely in accordance with the manufacturer’s protocol for Total Transcriptome Isolation except with a final ethanol concentration of 66% to increase the yield of smaller RNA species. Analysis of extracted RNA was performed following depletion of ribosomal RNA molecules using a commercial rRNA depletion kit for mixed bacterial samples (Lexogen, RiboCop META, #125). The ribo-depleted RNA samples were first fragmented using ultrasound (4 pulses of 30 s at 4°C). Then, an oligonucleotide adapter was ligated to the 3’ end of the RNA molecules. First-strand cDNA synthesis was performed using M-MLV reverse transcriptase with the 3’ adapter as primer. After purification, the 5’ Illumina TruSeq sequencing adapter was ligated to the 3’ end of the antisense cDNA. The resulting cDNA was PCR-amplified using a high-fidelity DNA polymerase and the barcoded TruSeq-libraries were pooled in approximately equimolar amounts. Sequencing of pooled libraries, spiked with PhiX control library, was performed at a minimum of 7 million reads per sample in single-ended mode with 100 cycles on the NextSeq 2000 platform (Illumina). Demultiplexed FASTQ files were generated with bcl-convert v4.2.4 (Illumina). Raw sequencing reads were subjected to quality and adapter trimming via Cutadapt (26) v2.5 using a cutoff Phred score of 20 and discarding reads without any remaining bases (parameters: --nextseq-trim=20 -m 1 -a AGATCGGAAGAGCACACGTCTGAACTCCAGTCAC). Afterwards, all reads longer than 11 nt were aligned to the *E. coli* K12 MG1655 reference genome (RefSeq assembly accession: GCF_000005845.2) using the pipeline READemption (27) v2.0.3 with segemehl version 0.3.4 (28) and an accuracy cut-off of 95% (parameters: -l 12 -a 95). READemption gene_quanti was applied to quantify aligned reads overlapping genomic features by at least 10 nt (-o 10) on the sense strand based on RefSeq annotations (CDS, ncRNA, rRNA, tRNA) for assembly GCF_000005845.2 from Mar 11, 2022. Based on these counts, differential expression analysis was conducted via DESeq2 (29) version 1.24.0. Read counts were normalized by DESeq2 and fold-change shrinkage was conducted by setting the parameter betaPrior to TRUE. Differential expression was assumed at adjusted p-value after Benjamini-Hochberg correction (padj) < 0.05 and |log2FoldChange| ≥ 1. Results of the DESeq2 analysis can be found in Supplementary Data 1.

### Immunoblotting

Immunoblotting was conducted in accordance to standard laboratory protocols, with primary antibodies incubated overnight at 4°C. The following antibodies were used: rabbit polyclonal anti-mCherry (abcam, ab167453), mouse monoclonal anti-DnaK antibody (Enzo, 8E2/2) was at 1:1000 dilution, HRP Goat anti-mouse IgG (BioLegend, 405306) at 1: 10, 000 dilution and HRP Goat anti-rabbit IgG (GE Healthcare, NA934) at 1:10,000 dilution. ECL Prime Western blotting detection reagent (GE Healthcare, RPN2232) was used to develop the blots, which were analysed on the ChemiDoc MP imaging system and bands quantified using Image Lab software.

## Results

### Hfq condensation is a response exclusively to intracellular changes

Our experimental system involves growing a batch culture of *E. coli* strain MG1655 (which contains photoactivatable mCherry tag fused to the C-terminal end of *hfq*) in a defined minimal growth medium in the presence of a limiting amount (3mM) of ammonium chloride as the sole N source and excess (22mM) glucose as the carbon (C) source (12). Under these conditions, when ammonium chloride in the growth medium runs out (N-), the bacteria enter a state of N starvation and become growth-arrested. However, sufficient glucose still remains to support growth if the N source becomes replenished (12). The Hfq-condensates are not detected at N-but form gradually as N starvation ensues and become detectable ∼6 hours into N starvation (N-6) and, by 24 hours following N-starvation (N-24) ∼80-90% of all cells in the population contain the Hfq-condensates. Further, Hfq-condensates also form in bacteria that have been treated (at N-) with rifampicin (transcription inhibitor) or chloramphenicol (translation inhibitor), implying that Hfq-condensates does not depend on *de novo* gene expression during N starvation (12). We considered whether Hfq condensation is triggered in response to an extracellular cue that accumulates over time under N starvation. To investigate this, we used bacteria from N+ (when cells are not N-starved, and Hfq-condensates are absent) and N-(when cells are N-starved, but Hfq-condensates are absent) and resuspended the bacteria in ‘spent’ media from cultures of bacteria grown to N-24. Although the spent media contains glucose (see above), it does not have any N source to support growth. We then periodically monitored the diffusion dynamics of individual Hfq molecules by photoactivated localisation microscopy (PALM) in live cells. We reasoned that if an extracellular cue existed, then Hfq-foci formation would occur faster in spent media (in which such a cue would have accumulated) than it would when bacteria become progressively more N-starved over time in normal (unspent) media. A quantitative parameter to measure Hfq-condensate formation, %H_IM_, was calculated as the proportion of total Hfq molecules with an apparent diffusion less than 0.08 um^2^/s, which had previously been defined as the ‘immobile’ population of Hfq molecules (12). The %H_IM_ was calculated based on all trajectories of Hfq in 50-200 bacterial cells within a given field of view. We did not detect any differences in %H_IM_ as a function of time following inoculation of bacteria from N+ (**Figure 1A**) or N-(**Figure 1B**) into spent media and, in both cases, the Hfq-condensates became detectable and clearly distinguishable from the mobile population of Hfq molecules ∼6 hours following incubation; the rate of Hfq condensation in both cases was similar to that observed in bacteria becoming progressively N-starved in unspent media (**Figure 1C**). We conclude that Hfq condensation is not mediated by any extracellular cues but represents a response exclusively to changes that occur inside *E. coli* cells during N-starvation.

**Figure 1.**
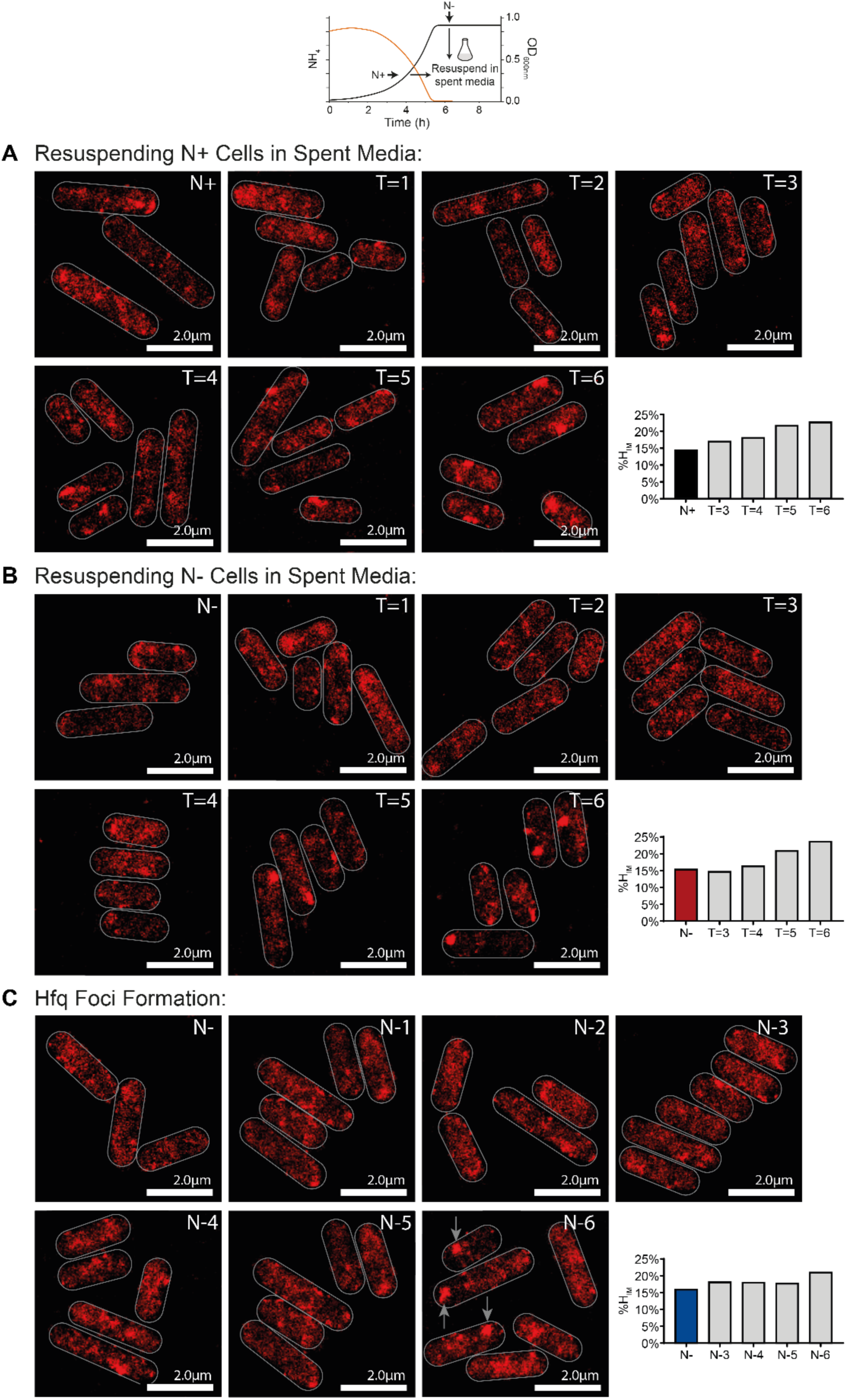
Hfq condensation is a response exclusively to intracellular changes. Representative PALM images of Hfq in *E. coli* after cells from **(A)** N+ or **(B)** N-were resuspended in spent media from N-24 cultures and imaged hourly. **(C)** *E. coli* cells experiencing N starvation for 6h, imaged hourly. Graphs show %HIM values.

### Cytoplasmic shrinkage during N starvation does not contribute to Hfq condensation

It is well established that the cytoplasm of *E. coli* cells shrinks during nutrient starvation (30,31). This inevitably increases the viscosity of the cytoplasm, affecting the diffusibility of macromolecules. We considered whether progressive shrinkage of the cytoplasm in response to N-starvation could trigger Hfq condensation. Therefore, we measured the average cytoplasmic area of bacterial cells and correlated this with the presence and dispersion of the Hfq-condensates. As shown in **Figure 2**, as expected, the average cytoplasmic area decreased from ∼1509 arbitrary units to ∼1235 arbitrary units when N+ bacteria became N-starved. Notably, no difference in the average cytoplasmic area was detected between N- (Hfq-condensates absent) and N-24 (Hfq-condensates present) bacteria. When N-24 bacteria, where the Hfq-condensates have fully formed, were resuspended in fresh media that supported growth recovery (ammonium chloride and glucose present; N+/C+) and led to the dispersion of the Hfq-condensates, the average cytoplasmic area reverted to that seen in N+ bacteria. However, when N-24 bacteria were resuspended in fresh media which only contained ammonium chloride but no glucose (N+/C-) and hence did not support growth recovery but caused the dispersion of the Hfq-condensates, the average cytoplasmic area did not increase and resembled that seen in N-24 bacteria (where the Hfq-condensates exist). Conversely, in control experiments when N-24 bacteria were resuspended in fresh media that contained no ammonium chloride but only glucose, where the Hfq-condensates *did not* disperse, the average cytoplasmic area also did not increase and resembled that seen in N-24-bacteria. Overall, we conclude that cytoplasmic shrinkage during N starvation does not contribute to Hfq condensation.

**Figure 2.**
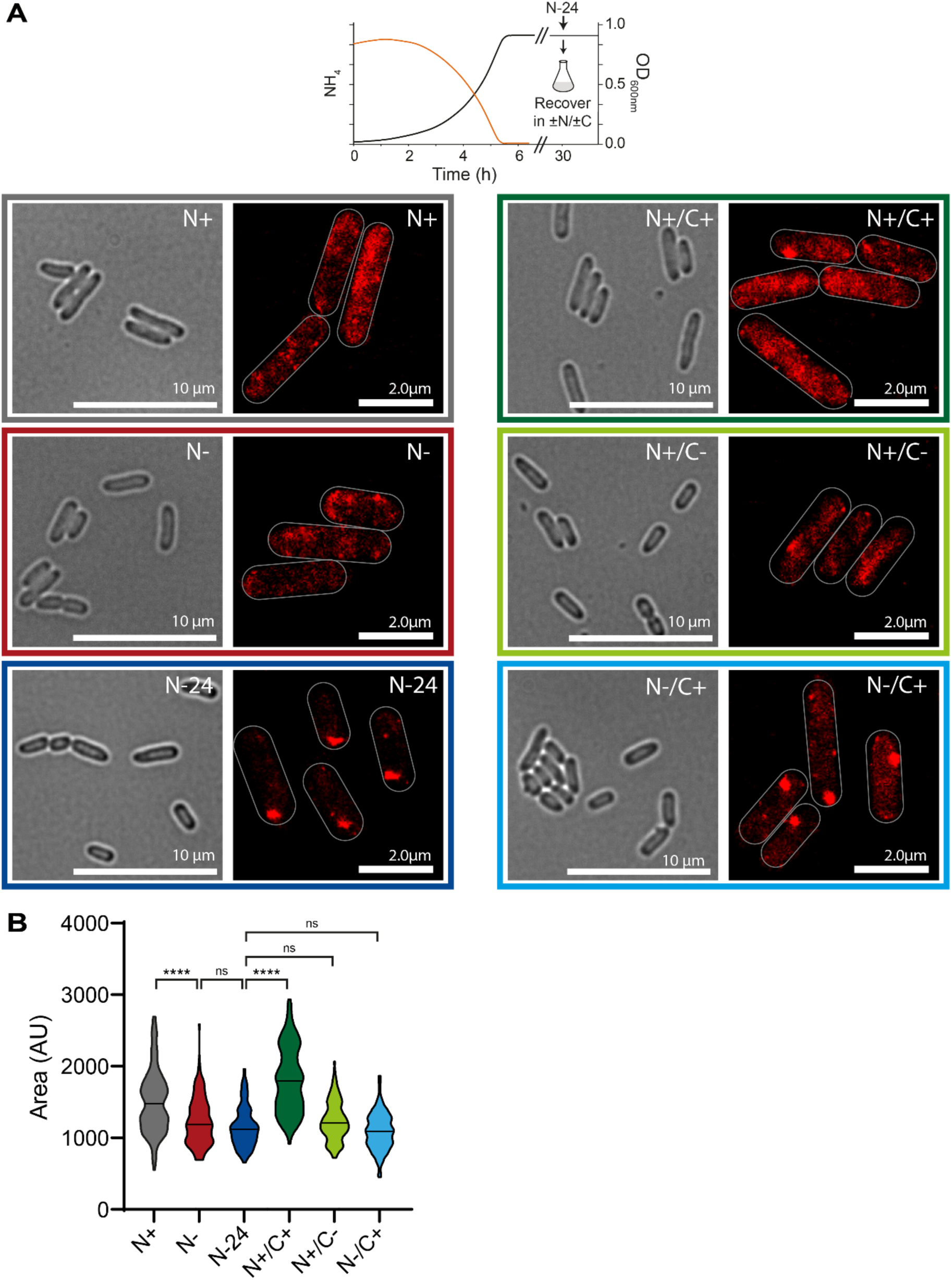
Cytoplasmic shrinkage during N starvation does not contribute to Hfq condensation. **(A)** Representative phase contrast of *E. coli* and PALM images of Hfq in *E. coli* at different stages of nitrogen starvation (N+, N- and N-24) and 2 hours after N-24 cells were resuspended in fresh media with different combinations of N and C (N+/C+, N+/C- and N-/C+). **(B)** Violin plot of bacterial cell area determined at each measured time point; the median area is indicated with the horizontal line. Statistical analysis performed by Kruskal-Wallis test with Dunn’s multiple comparisons. (****, P<0.0001).

### Hfq condensation in N-starved *E. coli* requires TmaR

Based on our results thus far, we focused our investigation on intracellular events that drive or require Hfq condensation. The dependency of Hfq-condensates on TmaR has been demonstrated in late stationary phase and osmotically stressed *E. coli* but not in N-starved *E. coli*. In addition, the Hfq-condensates and the TmaR-condensates colocalise in stationary phase and osmotically stressed *E. coli* and, notably, in both stress conditions, Hfq condensation in N-starved bacteria is dependent on TmaR (13). To investigate the role of the TmaR on Hfq condensation under N-starvation, we compared the diffusion dynamics of Hfq and TmaR (containing a photoactivatable mCherry tag fused to its C-terminal end) by PALM. As shown in **Figure 3A**, the percentage of cells containing the TmaR-condensates steadily increased during N starvation and, by N-24 ∼80% of cells contained the TmaR-foci (**Figure 3B**). In N-24 Δ*tmaR* bacteria we did not detect the Hfq-condensates (**Figure 3C**). We note that, unlike the Hfq-condensates, under our conditions, a substantial proportion of cells contain TmaR-condensates before the Hfq-foci become detectable. For example, at N+ and N-, when Hfq-condensates are entirely absent, ∼40% and ∼60% of the cells contained TmaR-condensates. It is known that glucose uptake rates in N-starved *E. coli* decreases more drastically than any other stress condition (32). As the TmaR condensation leads to the inhibition of PTS enzyme I (14,15), the results imply that the inhibition of glucose uptake in bacteria entering N starvation is not homogenous.

**Figure 3.**
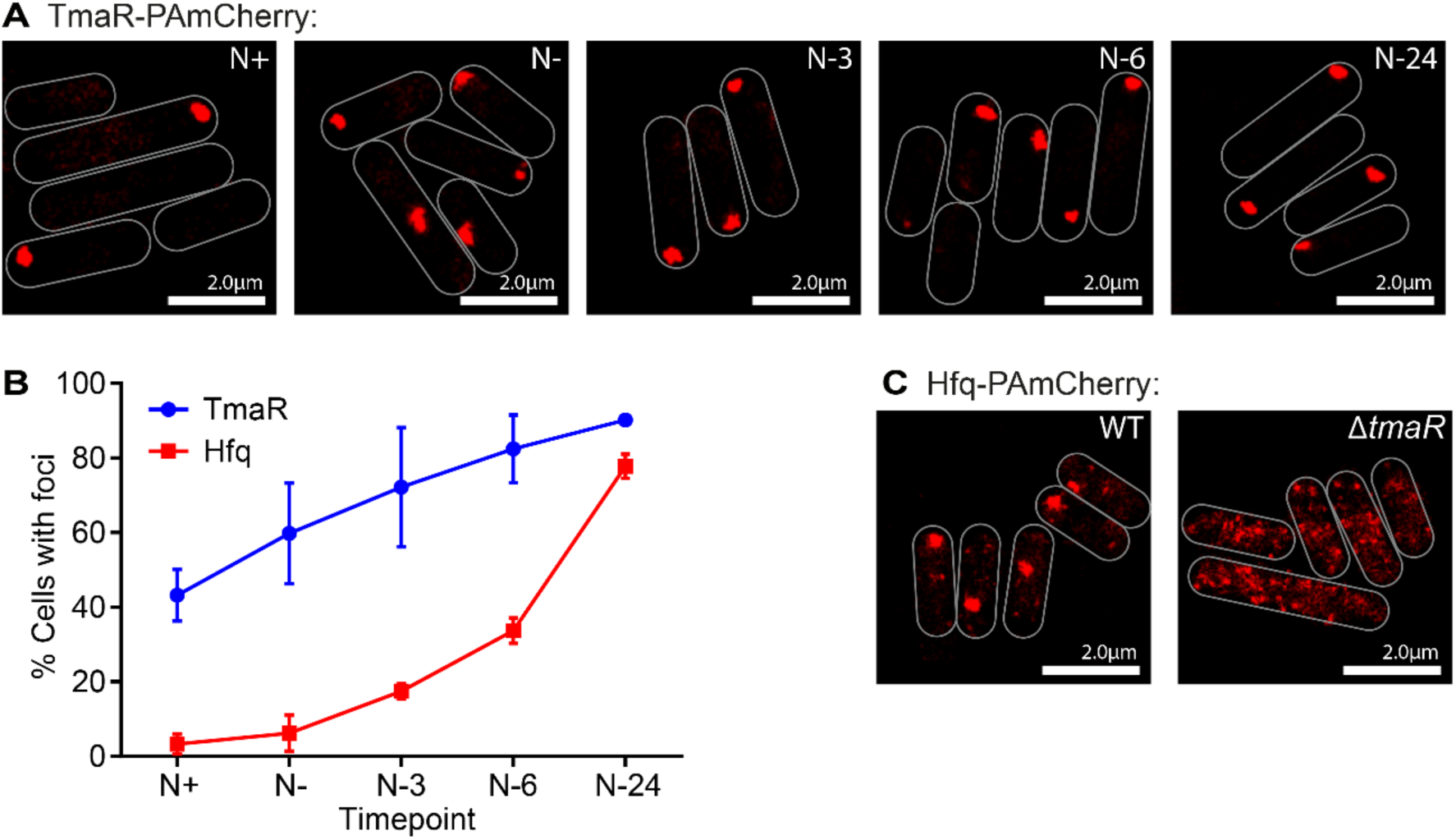
Hfq condensation in N-starved *E. coli* requires TmaR. **(A)** Representative PALM images of TmaR in *E. coli* as a function of time under N starvation. Images were taken at the indicated time points. **(B)** Graph showing the proportion of cells with detectable TmaR (blue) or Hfq (red) condensates during N starvation. Errors bars represent standard deviation (n>3). **(C)** Representative PALM images of Hfq in WT and Δ*tmaR* bacteria at N-24.

### Hfq condensation is an adaptative response that occurs independently of the canonical NtrBC pathway

Nitrogen starvation reduces the flux of α-ketoglutarate (αKG) out of the Krebs cycle into amino acids biosynthesis, leading to a rise in αKG levels, repression of glucose uptake and inhibition of cAMP synthesis. The latter prevents the global transcription factor CRP from activating transcription of genes required for uptake and catabolism of other sugars. Thus, αKG is a critical metabolite for coordinating the regulation of carbon assimilation during N starvation in *E. coli*. During growth in glucose, a particularly critical interaction linking N availability to glucose uptake is the inhibition of the PTS enzyme I, the target of TmaR (14,15), by αKG (33). As the dependency of Hfq condensation relies on TmaR, we speculated whether addition of exogenous αKG to N-bacteria, when Hfq-condensates are absent would cause Hfq condensation (recall that Hfq-condensates only become detectable by N-6). As shown in **Figure 4A and 4B**, when dimethyl-ketoglutarate (dmKG), a membrane-permeable ester that is cleaved by intracellular esterases to form αKG, was added to N-bacteria, Hfq condensation could be ‘induced’ and Hfq-condensates formed much sooner in dmKG treated compared to untreated bacteria. Control experiments showed that the dynamics of the TmaR foci remained unaffected by dmKG (**Figure 4A** and **Supplementary Figure 1A**). Additional control experiments showed that dmKG induction of Hfq-condensates was dependent on both TmaR (**Supplementary Figure 1B**) and polyP (**Supplementary Figure 1C**). Notably, Hfq-condensates could be induced with dmKG even in the presence of the transcription inhibitor rifampicin and translation inhibitor chloramphenicol, suggesting that the induction of Hfq condensation by dmKG occurs independently of any changes in gene expression due to the addition of dmKG (**Supplementary Figure 2**). Recall that the adaptive response to N starvation requires the two-component system NtrBC, where NtrB is the sensor kinase and NtrC the transcription regulator (see Introduction). At a low αKG concentration (e.g. under N-replete conditions), GlnB (PII) binds to αKG, which allows it to interact with NtrB, inhibiting its kinase activity and activating its phosphatase activity to dephosphorylate NtrC, rendering the latter unable to activate transcription. However, at higher αKG concentrations (e.g. upon N starvation), GlnB binds additional molecules of αKG and is unable to interact with NtrB, such that NtrB acts as a kinase to phosphorylate NtrC, allowing it to activate transcription of genes required for adaptation to N starvation (**Supplementary Figure 3A**). We thus considered whether dmKG induces Hfq condensation by overstimulating the NtrC regulon by binding to GlnB. However, Hfq-condensates could still be induced with dmKG in Δ*glnB* bacteria (**Supplementary Figure 3B and 3C**). Therefore, it seems that, during N starvation, the condensation of Hfq is a response that occurs independently of the canonical NtrBC-dependent adaptive response pathway. Indeed, in the absence of NtrC (Δ*glnG*), Hfq-condensates form faster than in WT bacteria, underscoring the view that Hfq condensation is not associated with the canonical NtrBC-mediated adaptive response (**Supplementary Figure 3D**). This view is further supported by the fact that Hfq condensation, in both dKG untreated (12) and treated (**Supplementary Figure 2**) bacteria occurs independently of *de novo* gene expression.

**Figure 4.**
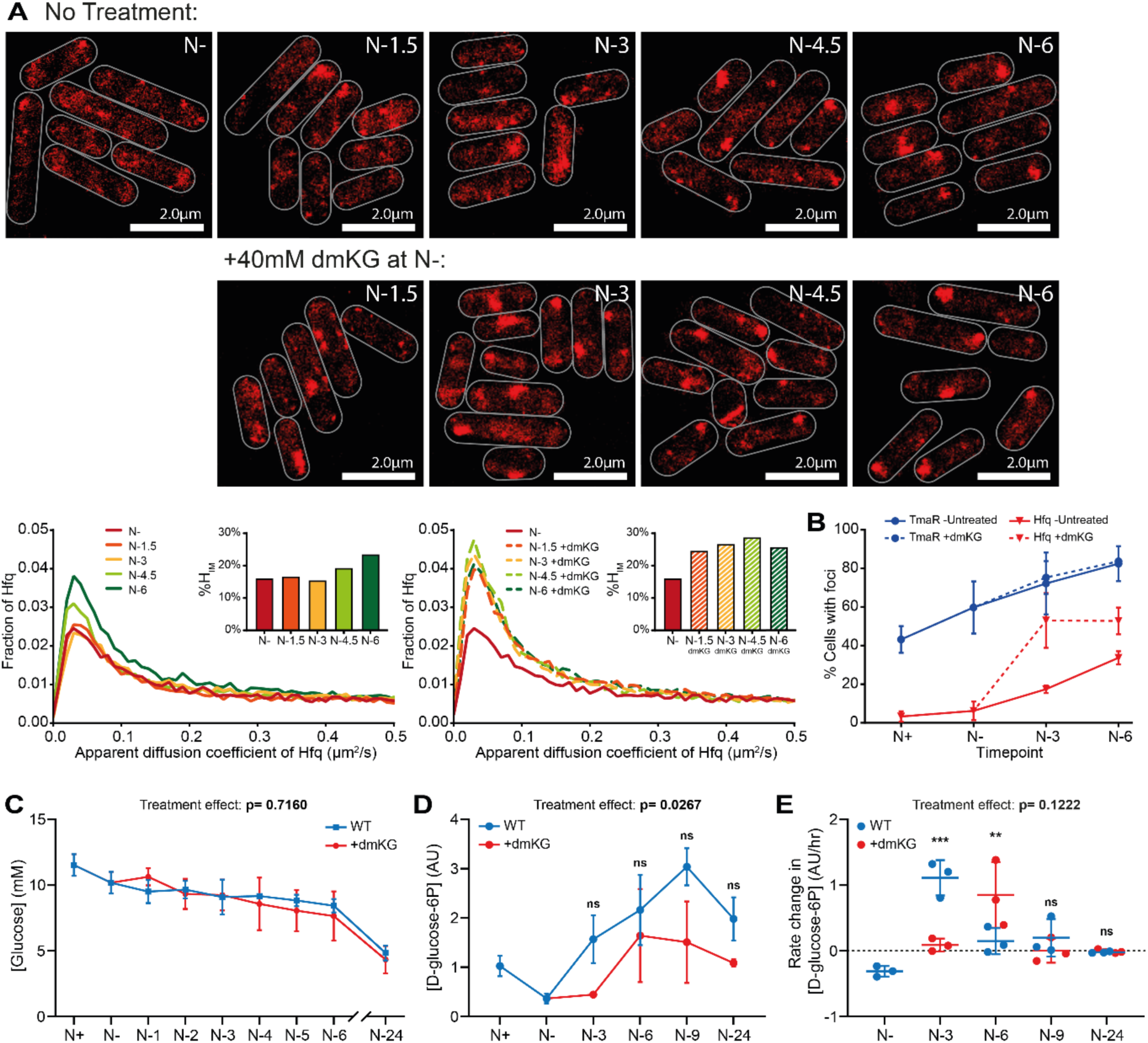
Hfq condensation in N-starved *E. coli* is coupled to inhibition of carbon assimilation. **(A)** Representative PALM images of Hfq in *E. coli* experiencing N starvation with and without treatment with 40mM dimethyl-ketoglutarate (dmKG) at N-. Graphs showing the distribution of apparent diffusion coefficient of Hfq molecules at the different sampling time points and the corresponding %H_IM_. **(B)** Graph showing the proportion of cells with detectable TmaR (blue) or Hfq (red) condensates during N starvation; the dashed line indicates the proportion of cells with detectable TmaR (blue) and Hfq (red) condensates during N starvation after adding dmKG at N-. Errors bars represent standard deviation (n>3). **(C)** Graph showing the concentration of glucose in the media in cultures of *E. coli* experiencing N starvation with (red) and without (blue) treatment with 40mM dmKG at N-. Errors bars represent standard deviation (n=3). **(D)** As in **(C)** but of intracellular concentration of D-glucose-6-phosphate (G6P). **(E)** Graph showing the rate change in G6P concentration; values are shown as a proportional range relative to the previous time point divided by time between time points (n=3). Statistical analysis performed by two-way ANOVA with Šidák multiple comparisons. (***, P<0.001; *, P<0.05).

## Hfq condensation in N-starved *E. coli* occurs concomitantly with the inhibition of glucose uptake

Given that: addition of dmKG induces the Hfq-condensates, αKG contributes to the inhibition of glucose uptake via inhibiting PTS enzyme I (see above) and Hfq-condensates can form independently of the NtrBC pathway, we considered whether Hfq condensation in N-starved *E. coli* occurs concomitantly with the inhibition of glucose uptake. To investigate whether the addition of dmKG indeed led to decreased glucose uptake, we measured extracellular glucose concentration as a function of time during N starvation from cultures of dmKG treated and untreated bacteria. As shown in **Figure 4C**, the change in extracellular glucose concentration as a function of time under N starvation did not substantially differ in the media of bacteria treated with dmKG and untreated controls (p=0.7160). However, we note that following the onset of N starvation (N-), the rate of decrease of glucose in the media (i.e. glucose uptake) of untreated bacteria was very low and therefore any differences between the treated and untreated samples would be too small to detect by measuring extracellular glucose concentrations. Hence, we used LC-MS to investigate the intracellular concentration of glucose-6-phosphate (G6P), the metabolic intermediate into which the majority of glucose is converted to during import into the cell. As shown in **Figure 4D**, in dmKG untreated bacteria, G6P concentration gradually increased during the first 9 hours of N starvation. In contrast, we observed a notable delay in the increase of G6P in the dmKG treated bacteria. Whilst this difference was not found to be significant at any individual time point, the overall treatment effect across all times points was found to be significant (p=0.0267). We next calculated the rate of change in G6P concentration between time points (calculated as per hour). As shown in **Figure 4E**, the rate of change in G6P concentration between N- (when dmKG was added) and N-3 was significantly lower in the dmKG treated cells than in untreated cells. Following N-3, rate of change in G6P concentration largely stabilised between dmKG treated and untreated cells, with rate of change in G6P concentration falling in the untreated cultures by N-6. We note that rate of change in G6P concentration represents the cumulative effect of changes in both uptake and metabolic use. Thus, the results must be interpreted with this consideration in mind. Nonetheless, the results clearly indicate an overall decrease in glucose uptake following dmKG treatment that concomitantly leads to Hfq condensation. We further note that Hfq condensation in the untreated (at N-6) and dmKG treated (at N-3) correlates with a period of decreased change in the concentration of G6P, and thus inhibition of glucose uptake during N starvation. We conclude that Hfq condensation in N-starved bacteria occurs concomitantly with the inhibition of glucose uptake.

### Hfq-condensates contribute to stabilising sRNAs in N-starved *E. coli*

We recently defined the Hfq-mediated RNA-RNA interactions during N starvation (10) and further demonstrated that the absence of Hfq specifically leads to the destabilisation of most Hfq-associated sRNA species in N-24 *E. coli* (34). Given that ∼50% of Hfq molecules are within the Hfq-condensates at N-24, we considered whether the Hfq-condensates have a role in sRNA stabilisation. As the Hfq-condensates do not form in Δ*tmaR* bacteria and therefore, unlike dmKG treatment, represents a binary response with respect to Hfq condensation, (i.e. Hfq-condensates present in wild-type bacteria and Hfq-condensates entirely absent in Δ*tmaR* bacteria), we compared the transcriptomes of N-24 wild-type and Δ*tmaR* bacteria to determine how the absence of TmaR, *ipso facto*, Hfq-condensates, affects sRNA abundance at N-24. We defined differentially expressed genes as those with expression levels changed ≥2-fold with a false discovery rate adjusted P < 0.05. According to this threshold, the absence of TmaR led to a dysregulation of ∼305 genes (∼89 and ∼216 genes up- and downregulated, respectively), corresponding to ∼7% of all genes in *E. coli* (**Figure 5A**). Put differently, despite the significant subcellular difference in WT and Δ*tmaR* bacteria (i.e. presence or absence of Hfq-condensates, respectively) the absence of TmaR only had a modest impact on the transcriptome of *E. coli* following 24h of N-starvation. Notably, although there is no detectable difference in Hfq levels in N-24 WT and Δ*tmaR* bacteria (**Supplementary Figure 4**), of the 25 sRNAs that interacted with Hfq at N-24 (10), 6 (24%) were downregulated in N-24 Δ*tmaR* bacteria, suggesting that the Hfq-condensates contributes to sRNA stabilisation in N-starved bacteria (**Figure 5B**). In further support of this, of 70 sRNAs and ncRNAs which *did not* interact with Hfq at N-24 (10), only 5 (7%) were downregulated in N-24 Δ*tmaR* bacteria (**Supplementary Figure 5**).

**Figure 5.**
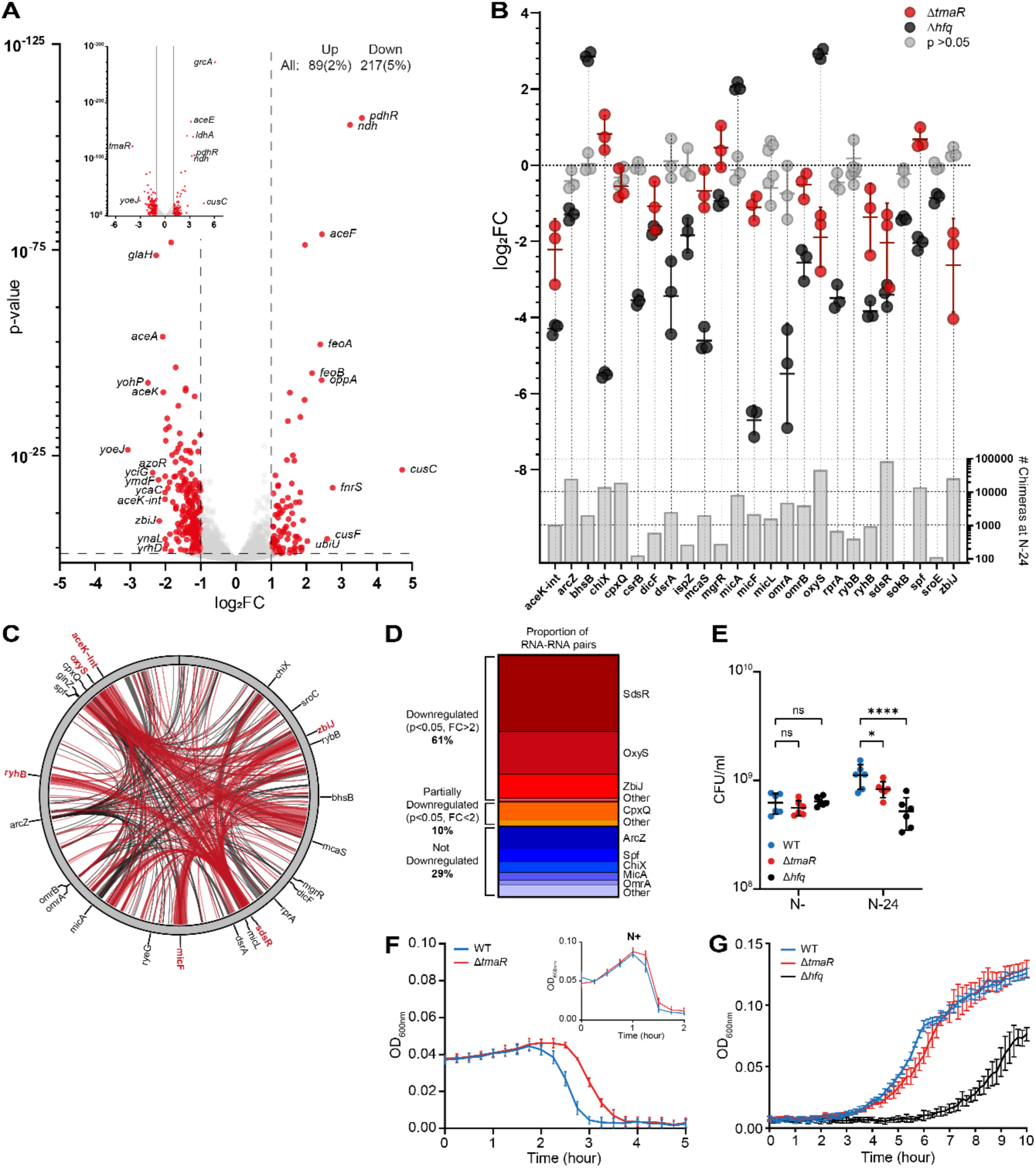
Hfq-condensates contribute to stabilising sRNAs in N-starved *E. coli*. **(A)** Volcano plots of differential RNA abundance in the transcriptome of Δ*tmaR E. coli* at N-24 as a log_2_ change from WT bacteria. Analysis performed by DESeq2. RNA differentially expressed more than 1 log_2_ (i.e. a greater than 2-fold change) are shown in blue. RNA differentially expressed more than 2 log_2_ (i.e., a greater than 4-fold change) are labelled. Inset was added to allow viewing of genes with very low p-values. The number and percentage (of total detected) of differentially expressed RNA are indicated at the top right. **(B)** Dot plot showing log_2_ fold change of individual sRNA known to interact with Hfq at N-24 in Δ*tmaR* (blue) and Δ*hfq* (red) bacteria relative to WT bacteria. The sRNAs that were not found to be differentially expressed with a p-value <0.05 by DESeq2 in each strain are shown in grey. Lower graph indicates the total number of chimeras found to contain the indicated sRNA at N-24 (across three replicates), as determined by RIL-seq (10). **(C)** Circos plot showing RNA-RNA pairs bound to the surface of Hfq in N-24 WT bacteria as previously determined by RIL-seq (10). Shown in red are interactions that involve sRNAs that are significantly downregulated (p<0.05, FC>2) in N-24 Δ*tmaR E. coli* at N-24 (as in (A) and (B)). The thickness of each connection is proportional to the average number of chimeras detected for a given interaction across three replicates. **(D)** Graph showing the proportion of the Hfq-bound RNA-RNA pairs, as previously determined by RIL-seq (10) which contain a specific sRNA, and whether those sRNA were downregulated in N-24 Δ*tmaR* bacteria. **(E)** Graph showing proportion of viable cells in the populations of Δ*tmaR* and Δ*hfq* bacteria following 20 min (N-) and 24 h (N-24) of N starvation. Error bars represent standard deviation (n = 6). Statistical analysis performed by two-way ANOVA with Dunnett’s multiple comparisons test. **(F)** Graph showing collapse of wild-type and Δ*tmaR* N-24 cultures following infection with T7 phage. Insert graph shows collapse of wild-type and Δ*tmaR* N-cultures following infection with T7 phage. Error bars represent standard deviation (*n* = 3). (*, P<0.05; ****, P<0.0001). **(G)** Growth-recovery of WT, Δ*tmaR* and Δ*hfq* N-24 bacteria following subculturing into fresh culture media. Error bars represent standard deviation (*n* = 3).

Strikingly, the sRNA that were downregulated in N-24 Δ*tmaR* bacteria contribute to ∼60% of detected RNA-RNA pairs bound to Hfq at N-24 (**Figure 5C and Figure 5D)** (10), supporting the notion that the formation of Hfq-condensates facilitates Hfq’s role as an sRNA stabiliser during N starvation. Therefore, we considered whether the absence of Hfq-condensates would perturb the RNA-RNA interaction network *ipso facto* post-transcriptional regulation at N-24, which consequently could compromise cellular homeostasis. In support of this view, the number of viable cells in the population of N-24 Δ*tmaR* bacteria were ∼25% less than in the N-24 WT population (**Figure 5E**). We note that this decrease in viable cells was not as great as that seen in the population of N-24 Δ*hfq* (∼53% decrease) bacteria (**Figure 5E**). The difference in viability between N-24 Δ*tmaR* and Δ*hfq* bacteria is perhaps unsurprising since the Hfq molecules not in the condensate in the Δ*tmaR* cells are still able to contribute to adaptive post-transcriptional regulation during N starvation. The efficacy by which bacteriophages infect and replicate in bacteria can serve as an indicator of bacterial metabolic ‘health’. Therefore, we measured how quickly the prototypical *E. coli* bacteriophage T7 replicated and caused collapse of the N-24 WT (Hfq-condensates present) and Δ*tmaR* (Hfq condensates absent) cultures. As shown in **Figure 5F**, following addition of T7, the culture of N-24 Δ*tmaR* bacteria collapsed ∼0.5h after that of the WT bacteria. Notably, this lag in culture collapse was not seen in with N+ bacteria where Hfq-condensates are absent in both the Δ*tmaR* and WT bacteria, confirming a role for Hfq-condensates in maintaining cellular homeostasis. However, whereas N-24 Δ*hfq* bacteria were also compromised for growth-recovery following inoculation into fresh growth media, Δ*tmaR* bacteria recovered like WT bacteria (**Figure 5G)**, underscoring a role for Hfq-condensates specifically in the cellular homeostasis during growth-arrest. Overall, we conclude that the Hfq-condensates contribute to preserving the stability of sRNAs during N starvation and thereby to cellular adaptation to N starvation.

## Discussion

Ribonucleoprotein-condensates are an emergent feature in the subcellular landscape of stressed bacteria and, like analogous condensates such as stress granules or P-bodies in eukaryotic cells (35,36), they contribute to the adaptive metabolism and homeostasis of RNA. In bacteria, key proteins involved in RNA synthesis, RNA degradation and RNA regulation have been described to undergo phase condensation. For example, condensates of *E. coli* RNA polymerase have been proposed to serve as hub to store non-transcribing RNA polymerase molecules during growth transitions (37). The *Caulobacter crescentus* RNase E undergoes condensation and forms a multiprotein complex called the BR-body that confers *C. crescentus* cells stress resistance (38,39). Although the primary function of BR-bodies is to facilitate RNA decay (38), the complex network of protein-protein interactions that underpin their formation suggests that BR-bodies could have multiple roles in RNA metabolism in *C. crescentus* (40). Further, RNases in several other bacteria, such as RNase Y in *Bacillus subtilis* (41) and RNase J in *Helicobacter pylori* (42) form condensates, contributing to RNA decay. Recently, the *E. coli* RNA chaperone CsrA was described to form condensates with the RNA-degradation machinery and regulate the expression of virulence genes (43).

The most ubiquitous bacterial RNA chaperone Hfq, the subject of this study, forms polar condensates in response to diverse stresses in *E. coli* (see Introduction). Despite the prevalence of ribonucleoprotein condensates in stressed bacteria, most studies to date have focused on understanding their function in stress physiology through determining their composition. We have now shown that Hfq condensation is an adaptive response to N starvation. Inspired by the observation made by the Amster-Choder laboratory revealing that Hfq condensates co-localise with condensates of a novel glucose uptake regulator TmaR (13), we have now shown that the formation of Hfq-condensates occurs concomitantly with inhibition of glucose uptake in N-starved *E. coli*. An important breakthrough in our study is the finding that Hfq-condensates could be induced by αKG – the key metabolite involved in nitrogen and carbon metabolism in *E. coli*. It is well established that intracellular concentration of αKG increases upon entry into N starvation but is unlikely to rise further during long-term N starvation as it is found in the supernatants of N-starved cultures (suggesting active export of αKG to prevent its hyperaccumulation inside cells)(44). Thus, we propose that Hfq condensation is unlikely to be due to the accumulation of αKG as a function of time under N starvation. Interestingly, computational predictions indicated a potential interaction between Hfq and αKG (our unpublished observations). However, these predictions are not supported by experimental data from isothermal titration calorimetry and dynamic light scattering, which demonstrated that αKG does not bind to Hfq (our unpublished data). Therefore, the effect of αKG on Hfq is likely to be indirect and independent of TmaR, as TmaR-condensates were not detectably affected by αKG. Nonetheless, our results showing that Hfq condensation can be induced by αKG, which concomitantly results in the inhibition of glucose uptake, clearly indicates that Hfq condensation is a cellular process that is coupled to the inhibition of carbon assimilation during N starvation.

The majority of bacterial cells contain Hfq-condensates following 24h under N starvation. Therefore, we regard Hfq-condensates that exist in N-24 bacteria as fully formed. By leveraging the finding that Hfq-condensates do not form in N-24 Δ*tmaR* bacteria and our previous data describing the transcriptome (34) and genome-wide Hfq-associated RNA-RNA interaction network (10) at N-24, we have now revealed a function for Hfq-condensates in preserving sRNA stability in N-starved *E. coli*. We note that a limitation of this conclusion is the absence of evidence showing that the destabilized sRNA species in N-24 Δ*tmaR* bacteria are indeed physically associated with the Hfq-condensates, but this will be subject of future studies. Nonetheless, our conclusions are consistent with and expand results of a recent study from the Jakob laboratory showing Hfq-condensates that form under N starvation function to preserve polyadenylated mRNAs (which would otherwise be degraded) and mRNA encoding components of the translation machinery (17). We previously reported that condensates of the RNA degradosome colocalise with Hfq-condensates in N-starved *E. coli* (11). Therefore, in light of our new results, we propose that condensates of Hfq and the RNA degradosome interact to coordinate the selective preservation and degradation of RNA molecules during N starvation.

The bacterial transcriptional response to N-starvation is well-understood and is primarily triggered by the NtrBC system in *E. coli* and related bacteria (4). At the post-transcriptional level, the Hfq-mediated RNA-RNA interaction network undergoes large-scale changes upon entry into and during N starvation, underscoring the importance of the adaptive post-transcriptional response to N starvation (10). Pertinent to this study, the abundance of the Hfq-dependent sRNA GlnZ increases upon entry into N starvation (9). One of the targets of GlnZ is *sucA*, which encodes a subunit of αKG-dehydrogenase, the enzyme which catalyses the conversation of αKG to succinyl-CoA in the Krebs cycle. As GlnZ downregulates the expression of *sucA*, in light of our new results, we propose that GlnZ redirects αKG from the Krebs cycle to trigger Hfq condensation and concomitantly the inhibition of carbon assimilation and stabilisation of Hfq-associated sRNAs. We note that GlnZ levels do not differ between N-24 WT and Δ*tmaR* bacteria (**Supplementary Figure 6**). Interestingly, GlnZ also targets *tmaR* but the regulatory consequence of this interaction on *tmaR* expression is presently unclear. Nonetheless, our study now highlights that the post-transcriptional basis of the adaptive response to N starvation extends beyond canonical sRNA-mediated regulation and involves condensation of Hfq, stabilisation of sRNAs by Hfq-condensates and the inhibition of carbon assimilation and flux through the Krebs cycle.

In sum, our study has shown that Hfq-condensates are associated with the inhibition of carbon assimilation in N-starved *E. coli* and has expanded the current perception in the field that Hfq-condensates contribute to just preserving RNA molecules (17). Although both of these functions of Hfq-condensates might not be mutually inclusive, Hfq-condensates clearly contribute to maintaining cellular homeostasis during N starvation. Most bacterial stress ribonucleoprotein-condensates, including Hfq-condensates, are heterotypic in nature and their formation is underpinned by multivalent interactions between different proteins and different types of RNA molecules. Although we are now beginning to understand why they form and how they contribute to stress adaption, their heterotypic nature implies that neither their formation is triggered by a universal ‘stress signal’ nor they are restricted in their functions. Instead, as bacteria must respond and adapt to and survive ever-changing conditions, it is likely that the formation of ribonucleoprotein-condensates is precipitated by a combination of diverse cellular and physical factors and contribute to different adaptive, albeit RNA metabolism associated, functions in the cell.

## Acknowledgements

We thank Ben Luisi, Cambridge University, for help with the Hfq-αKG interaction assays and Martin Buck, Imperial College, for critical comments on manuscript.

## Supplementary data

Supplementary data is available at NAR online.

## Conflict of interest

None declared.

## Funding

This work was supported by the BBSRC (BB*/*V000284*/*1) and Leverhulme Trust (RPG-2020-050) project grants to S.W. H. E. was funded by an MRC ICASE PhD studentship. Funding to pay the Open Access publication charges for this article was provided by Imperial Open Access Fund.

## Data availability

The RNA-seq discussed in this publication is accessible through ArrayExpress: E-MTAB-14999.

---

**Supplementary Figure 1.**
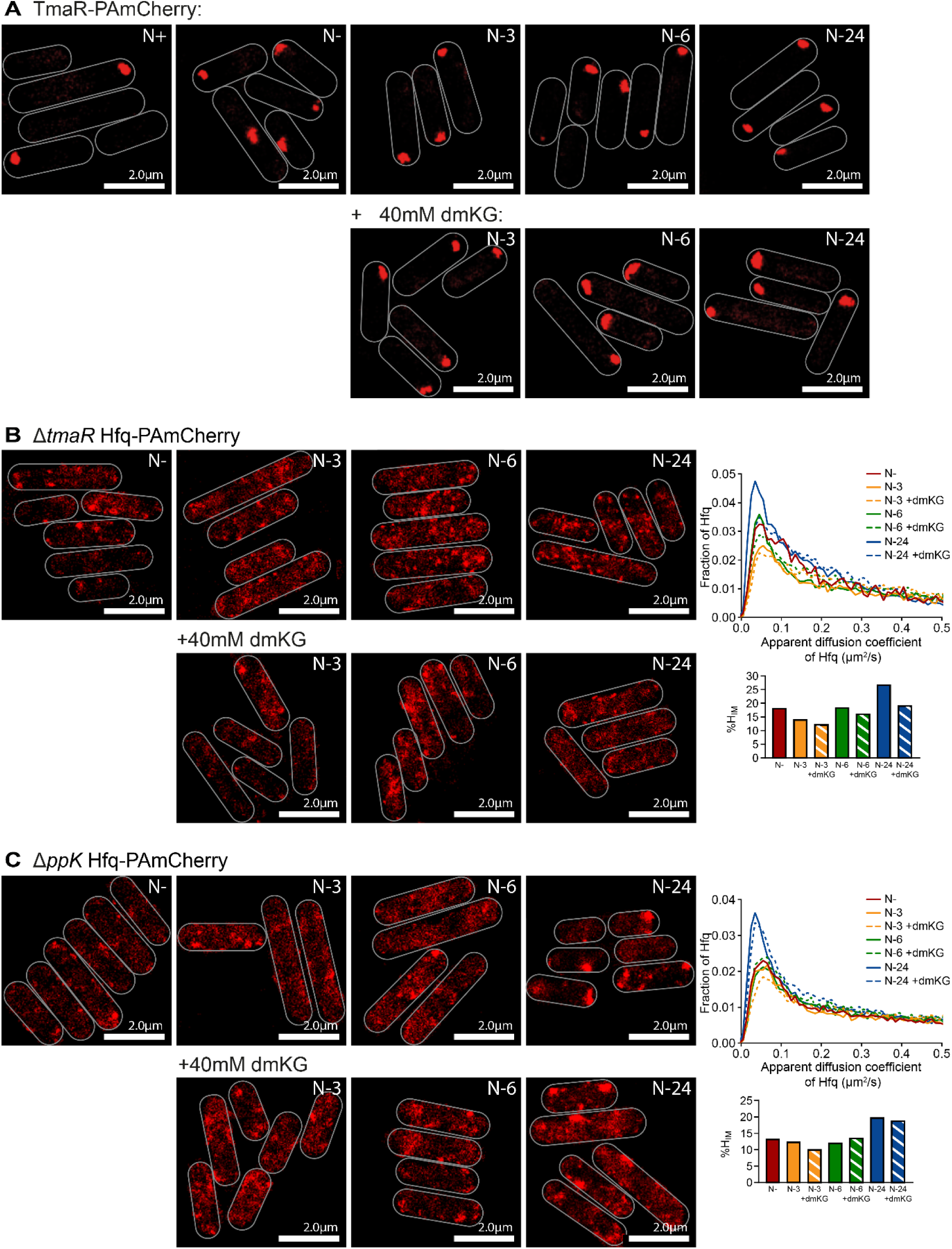
**(A)** Representative PALM images of TmaR in *E. coli* experiencing N starvation with and without 40mM dmKG added at N-. **(B)** As in (A) but of Hfq in Δ*tmaR* bacteria. (C) As in B but in Δ*ppK* bacteria. In (B) and (C), the graphs show the distribution of apparent diffusion coefficient of Hfq molecules at the different sampling time points and the corresponding %H_IM_.

**Supplementary Figure 2.**
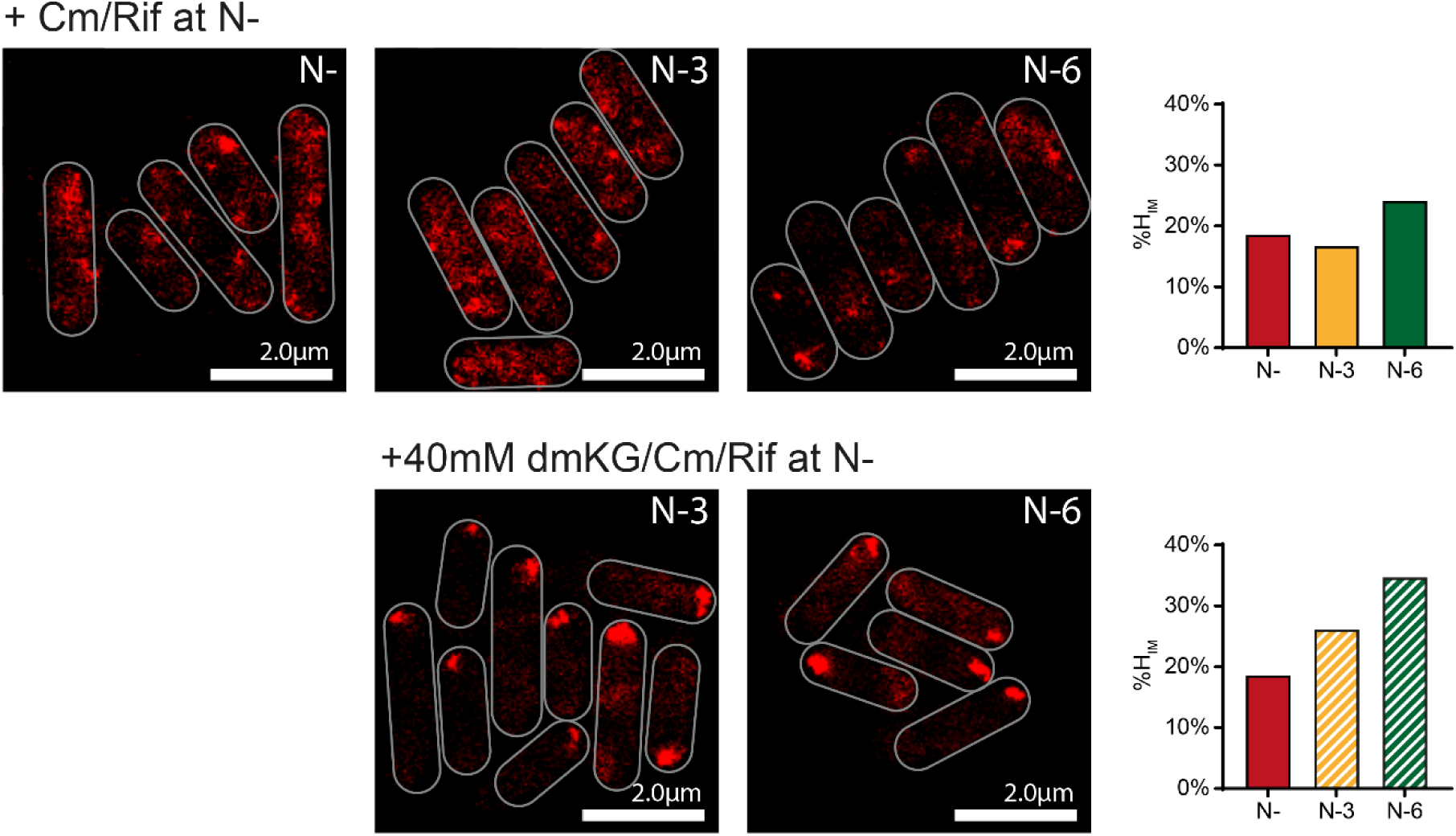
Representative PALM images of Hfq in *E. coli* experiencing N starvation, treated with 100μg/ml rifampicin and 150μg/ml chloramphenicol at N-, with and without treatment with 40mM dimethyl-ketoglutarate (dmKG) at N-. Graphs show %H_IM_.

**Supplementary Figure 3.**
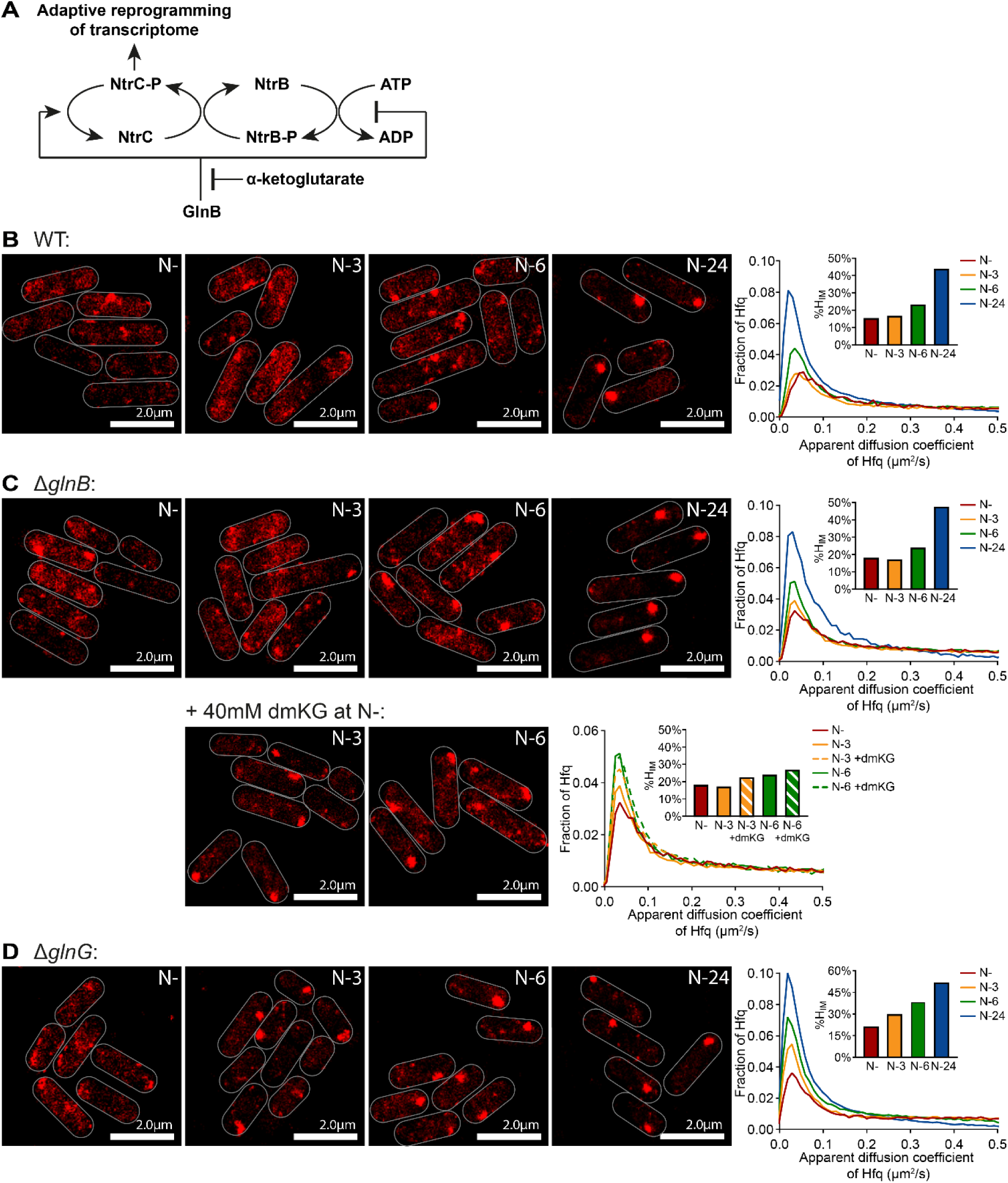
**(A)** Schematic showing GlnB and αKG mediated regulation of the NtrBC two-component system during nitrogen starvation. **(B)** Representative PALM images of Hfq in *E. coli* experiencing N starvation. Graphs showing the distribution of apparent diffusion coefficient of Hfq molecules at the different sampling time points and the corresponding %H_IM_. **(C)** as in **(B)** but for Δ*glnB* bacteria, with and without treated with 40mM dimethyl-ketoglutarate (dmKG) at N-. **(D)** as in **(B)** but for Δ*glnG* bacteria.

**Supplementary Figure 4.**
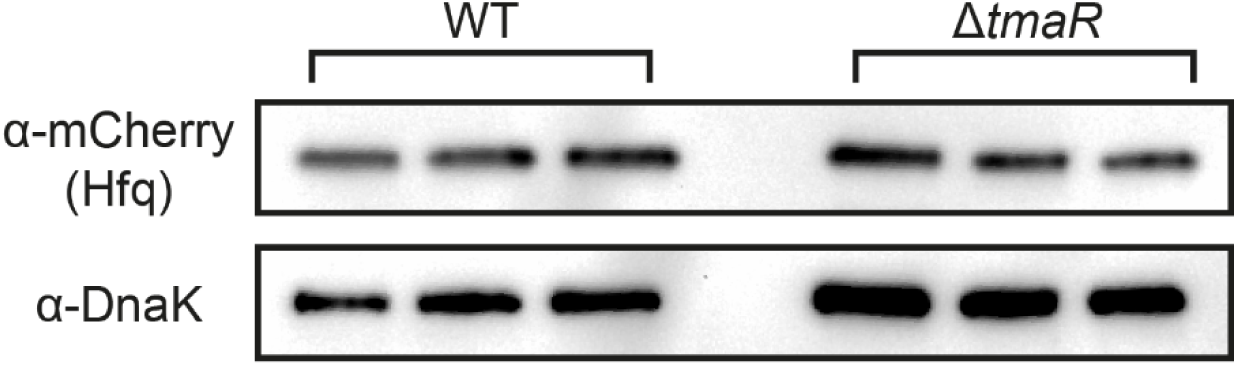
Immunoblot of whole-cell extracts of wild-type and Δ*tmaR E. coli* containing Hfq translationally fused to PAmCherry, sampled at N-24, three biological replicates are shown. Probed with anti-mCherry antibody (for Hfq) and anti-DnaK antibody (loading control).

**Supplementary Figure 5.**
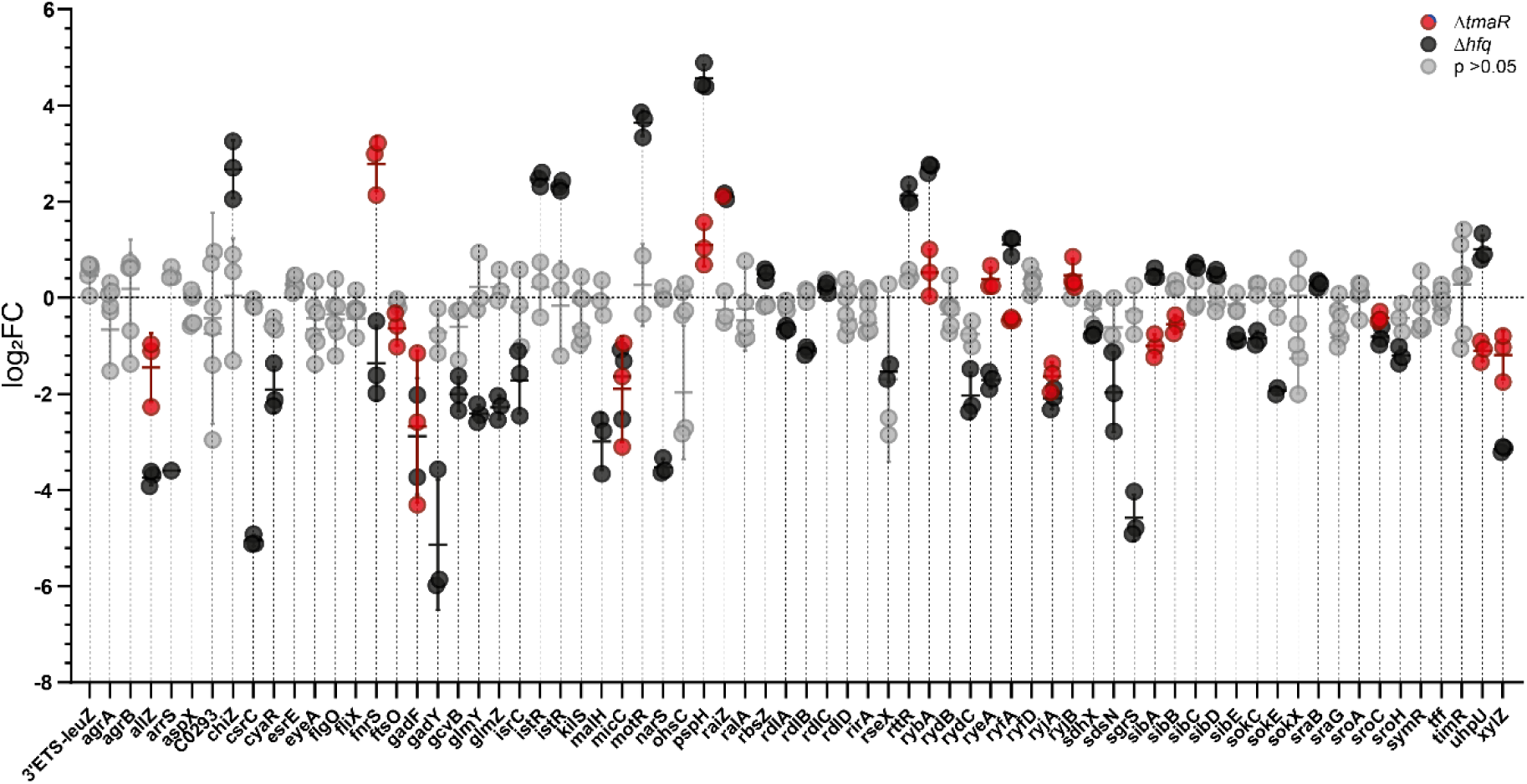
Dot plot showing log_2_ fold change of individual sRNA and non-coding RNA previously shown not to interact with Hfq at N-24 in Δ*tmaR* (red) and Δ*hfq* (black) bacteria relative to WT bacteria. Where an RNA was not found to be differentially expressed with a p-value <0.05 by DESeq2 in each strain are shown in grey.

**Supplementary Figure 6.**
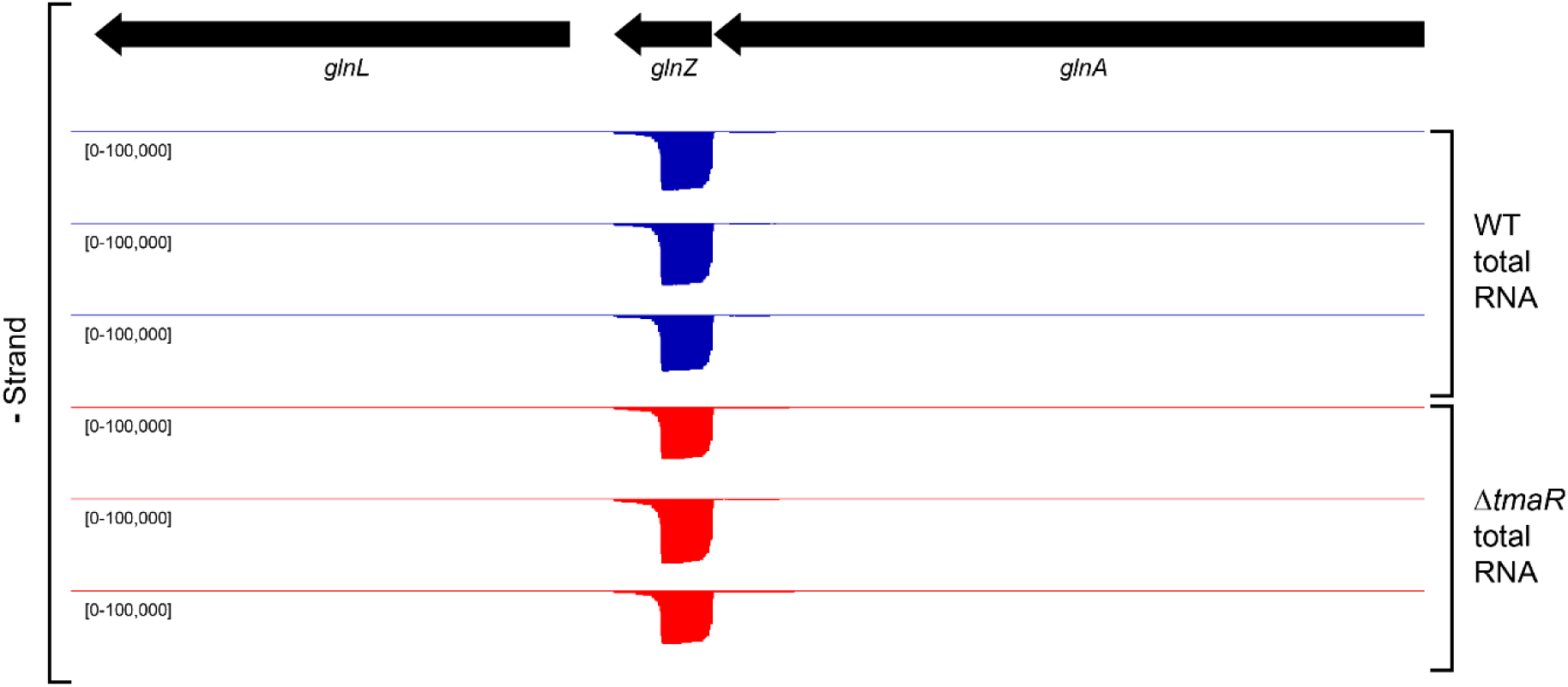
In browser images of normalised RNA reads mapped to the negative strand of the *glnA-glnZ* gene locus, from WT (blue) and Δ*tmaR* (red) bacteria grown to N-25 as determined by RNA-seq. Three replicates are shown, with identical scales.

## Notes

### Competing Interest Statement

The authors have declared no competing interest.

## References

1. Updegrove, T.B., Zhang, A. and Storz, G. (2016) Hfq: the flexible RNA matchmaker. Curr Opin Microbiol, 30, 133–138.

2. Vogel, J. and Luisi, B.F. (2011) Hfq and its constellation of RNA. Nat Rev Microbiol, 9, 578–589.

3. Papenfort, K. and Melamed, S. (2023) Small RNAs, Large Networks: Posttranscriptional Regulons in Gram-Negative Bacteria. Annu Rev Microbiol, 77, 23–43.

4. Reitzer, L. (2003) Nitrogen assimilation and global regulation in Escherichia coli. Annu Rev Microbiol, 57, 155–176.

5. Brown, D.R., Barton, G., Pan, Z., Buck, M. and Wigneshweraraj, S. (2014) Nitrogen stress response and stringent response are coupled in Escherichia coli. Nature communications, 5, 4115.

6. Aquino, P., Honda, B., Jaini, S., Lyubetskaya, A., Hosur, K., Chiu, J.G., Ekladious, I., Hu, D., Jin, L., Sayeg, M.K. et al. (2017) Coordinated regulation of acid resistance in Escherichia coli. BMC Syst Biol, 11, 1.

7. Zimmer, D.P., Soupene, E., Lee, H.L., Wendisch, V.F., Khodursky, A.B., Peter, B.J., Bender, R.A. and Kustu, S. (2000) Nitrogen regulatory protein C-controlled genes of Escherichia coli: scavenging as a defense against nitrogen limitation. Proceedings of the National Academy of Sciences of the United States of America, 97, 14674–14679.

8. Switzer, A., Burchell, L., McQuail, J. and Wigneshweraraj, S. (2020) The Adaptive Response to Long-Term Nitrogen Starvation in Escherichia coli Requires the Breakdown of Allantoin. Journal of bacteriology, 202.

9. Walling, L.R., Kouse, A.B., Shabalina, S.A., Zhang, H. and Storz, G. (2022) A 3’ UTR-derived small RNA connecting nitrogen and carbon metabolism in enteric bacteria. Nucleic Acids Res, 50, 10093–10109.

10. McQuail, J., Matera, G., Grafenhan, T., Bischler, T., Haberkant, P., Stein, F., Vogel, J. and Wigneshweraraj, S. (2024) Global Hfq-mediated RNA interactome of nitrogen starved Escherichia coli uncovers a conserved post-transcriptional regulatory axis required for optimal growth recovery. Nucleic Acids Res, 52, 2323–2339.

11. McQuail, J., Carpousis, A.J. and Wigneshweraraj, S. (2022) The association between Hfq and RNase E in long-term nitrogen-starved Escherichia coli. Molecular microbiology, 117, 54–66.

12. McQuail, J., Switzer, A., Burchell, L. and Wigneshweraraj, S. (2020) The RNA-binding protein Hfq assembles into foci-like structures in nitrogen starved Escherichia coli. The Journal of biological chemistry, 295, 12355–12367.

13. Goldberger, O., Szoke, T., Nussbaum-Shochat, A. and Amster-Choder, O. (2022) Heterotypic phase separation of Hfq is linked to its roles as an RNA chaperone. Cell Rep, 41, 111881.

14. Szoke, T., Albocher, N., Govindarajan, S., Nussbaum-Shochat, A. and Amster-Choder, O. (2021) Tyrosine phosphorylation-dependent localization of TmaR that controls activity of a major bacterial sugar regulator by polar sequestration. Proceedings of the National Academy of Sciences of the United States of America, 118.

15. Szoke, T., Goldberger, O., Albocher-Kedem, N., Barsheshet, M., Dezorella, N., Nussbaum-Shochat, A., Wiener, R., Schuldiner, M. and Amster-Choder, O. (2023) Regulation of major bacterial survival strategies by transcripts sequestration in a membraneless organelle. Cell Rep, 42, 113393.

16. Kannaiah, S., Livny, J. and Amster-Choder, O. (2019) Spatiotemporal Organization of the E. coli Transcriptome: Translation Independence and Engagement in Regulation. Molecular cell, 76, 574–589 e577.

17. Guan, J., Hurto, R.L., Rai, A., Azaldegui, C.A., Ortiz-Rodriguez, L.A., Biteen, J.S., Freddolino, L. and Jakob, U. (2025) HP-Bodies - Ancestral Condensates that Regulate RNA Turnover and Protein Translation in Bacteria. bioRxiv.

18. Rathnayaka-Mudiyanselage, I.W., Nandana, V. and Schrader, J.M. (2024) Proteomic composition of eukaryotic and bacterial RNA decay condensates suggests convergent evolution. Curr Opin Microbiol, 79, 102467.

19. Datsenko, K.A. and Wanner, B.L. (2000) One-step inactivation of chromosomal genes in Escherichia coli K-12 using PCR products. Proceedings of the National Academy of Sciences of the United States of America, 97, 6640–6645.

20. Baba, T., Ara, T., Hasegawa, M., Takai, Y., Okumura, Y., Baba, M., Datsenko, K.A., Tomita, M., Wanner, B.L. and Mori, H. (2006) Construction of Escherichia coli K-12 in-frame, single-gene knockout mutants: the Keio collection. Mol Syst Biol, 2, 2006 0008.

21. Atlas, R.M. ((2010)) Handbook of Microbiological Media (4th ed.). CRC Press.

22. Stracy, M., Lesterlin, C., Garza de Leon, F., Uphoff, S., Zawadzki, P. and Kapanidis, A.N. (2015) Live-cell superresolution microscopy reveals the organization of RNA polymerase in the bacterial nucleoid. Proceedings of the National Academy of Sciences of the United States of America, 112, E4390–4399.

23. Endesfelder, U., Finan, K., Holden, S.J., Cook, P.R., Kapanidis, A.N. and Heilemann, M. (2013) Multiscale spatial organization of RNA polymerase in Escherichia coli. Biophysical journal, 105, 172–181.

24. Behrends, V., Tredwell, G.D. and Bundy, J.G. (2011) A software complement to AMDIS for processing GC-MS metabolomic data. Anal Biochem, 415, 206–208.

25. Stead, M.B., Agrawal, A., Bowden, K.E., Nasir, R., Mohanty, B.K., Meagher, R.B. and Kushner, S.R. (2012) RNAsnap: a rapid, quantitative and inexpensive, method for isolating total RNA from bacteria. Nucleic acids research, 40, e156.

26. Martin, M. (2011) Cutadapt removes adapter sequences from high-throughput sequencing reads. EMBnet.journal, 17.

27. Forstner, K.U., Vogel, J. and Sharma, C.M. (2014) READemption-a tool for the computational analysis of deep-sequencing-based transcriptome data. Bioinformatics, 30, 3421–3423.

28. Hoffmann, S., Otto, C., Kurtz, S., Sharma, C.M., Khaitovich, P., Vogel, J., Stadler, P.F. and Hackermuller, J. (2009) Fast mapping of short sequences with mismatches, insertions and deletions using index structures. PLoS Comput Biol, 5, e1000502.

29. Love, M.I., Huber, W. and Anders, S. (2014) Moderated estimation of fold change and dispersion for RNA-seq data with DESeq2. Genome Biol, 15, 550.

30. Lange, R. and Hengge-Aronis, R. (1991) Growth phase-regulated expression of bolA and morphology of stationary-phase Escherichia coli cells are controlled by the novel sigma factor sigma S. Journal of bacteriology, 173, 4474–4481.

31. Parry, B.R., Surovtsev, I.V., Cabeen, M.T., O’Hern, C.S., Dufresne, E.R. and Jacobs-Wagner, C. (2014) The bacterial cytoplasm has glass-like properties and is fluidized by metabolic activity. Cell, 156, 183–194.

32. Chubukov, V. and Sauer, U. (2014) Environmental dependence of stationary-phase metabolism in Bacillus subtilis and Escherichia coli. Applied and environmental microbiology, 80, 2901–2909.

33. Doucette, C.D., Schwab, D.J., Wingreen, N.S. and Rabinowitz, J.D. (2011) alpha-Ketoglutarate coordinates carbon and nitrogen utilization via enzyme I inhibition. Nat Chem Biol, 7, 894–901.

34. McQuail, J., Krepl, M., Katsuya-Gaviria, K., Tabib-Salazar, A., Burchell, L., Bischler, T., Grafenhan, T., Brear, P., Sponer, J., Luisi, B.F. et al. (2025) Transcriptome-scale analysis uncovers conserved residues in the hydrophobic core of the bacterial RNA chaperone Hfq required for small regulatory RNA stability. Nucleic acids research, 53.

35. Banani, S.F., Lee, H.O., Hyman, A.A. and Rosen, M.K. (2017) Biomolecular condensates: organizers of cellular biochemistry. Nature reviews. Molecular cell biology, 18, 285–298.

36. Riggs, C.L., Kedersha, N., Ivanov, P. and Anderson, P. (2020) Mammalian stress granules and P bodies at a glance. J Cell Sci, 133.

37. Ladouceur, A.M., Parmar, B.S., Biedzinski, S., Wall, J., Tope, S.G., Cohn, D., Kim, A., Soubry, N., Reyes-Lamothe, R. and Weber, S.C. (2020) Clusters of bacterial RNA polymerase are biomolecular condensates that assemble through liquid-liquid phase separation. Proceedings of the National Academy of Sciences of the United States of America, 117, 18540–18549.

38. Al-Husini, N., Tomares, D.T., Pfaffenberger, Z.J., Muthunayake, N.S., Samad, M.A., Zuo, T., Bitar, O., Aretakis, J.R., Bharmal, M.M., Gega, A. et al. (2020) BR-Bodies Provide Selectively Permeable Condensates that Stimulate mRNA Decay and Prevent Release of Decay Intermediates. Molecular cell, 78, 670–682 e678.

39. Al-Husini, N., Tomares, D.T., Bitar, O., Childers, W.S. and Schrader, J.M. (2018) alpha-Proteobacterial RNA Degradosomes Assemble Liquid-Liquid Phase-Separated RNP Bodies. Molecular cell, 71, 1027–1039 e1014.

40. Nandana, V., Rathnayaka-Mudiyanselage, I.W., Muthunayake, N.S., Hatami, A., Mousseau, C.B., Ortiz-Rodriguez, L.A., Vaishnav, J., Collins, M., Gega, A., Mallikaarachchi, K.S. et al. (2023) The BR-body proteome contains a complex network of protein-protein and protein-RNA interactions. Cell Rep, 42, 113229.

41. Hamouche, L., Billaudeau, C., Rocca, A., Chastanet, A., Ngo, S., Laalami, S. and Putzer, H. (2020) Dynamic Membrane Localization of RNase Y in Bacillus subtilis. mBio, 11.

42. Tejada-Arranz, A., Galtier, E., El Mortaji, L., Turlin, E., Ershov, D. and De Reuse, H. (2020) The RNase J-Based RNA Degradosome Is Compartmentalized in the Gastric Pathogen Helicobacter pylori. mBio, 11.

43. Aroeti, L., Elbaz, N., Faigenbaum-Romm, R., Yakovian, O., Altuvia, Y., Argaman, L., Katsowich, N., Bejerano-Sagie, M., Ravins, M., Margalit, H. et al. (2025) Formation of a membraneless compartment regulates bacterial virulence. Nature communications, 16, 3834.

44. Schumacher, J., Behrends, V., Pan, Z., Brown, D.R., Heydenreich, F., Lewis, M.R., Bennett, M.H., Razzaghi, B., Komorowski, M., Barahona, M. et al. (2013) Nitrogen and carbon status are integrated at the transcriptional level by the nitrogen regulator NtrC in vivo. mBio, 4, e00881–00813.

